# The effects of fasting on ischemic infarcts in the rat

**DOI:** 10.1101/2022.11.15.516543

**Authors:** Anna M Schneider, Alastair M Buchan, Yvonne Couch

## Abstract

**Background:** Inflammation has been found to be largely detrimental early in the acute phase of stroke but beneficial at more chronic stages. Fasting has been shown to reduce inflammation acutely. We aimed to determine whether post-ischemic fasting improves stroke outcomes through attenuated inflammation.

**Methods:** After an endothelin-1 lesion was created in the striatum, animals were subjected to either normal feeding or water-only fasting for 24 hours.

**Results:** It was found that at 24 hours, fasting reduced infarct volume and BBB breakdown and lowered both circulating and brain neutrophils.

**Conclusions:** These findings suggest that fasting is a potentially beneficial non-pharmacological additive therapeutic option for cerebral ischemia, which might act by reducing inflammation in the acute disease stage.

## Introduction

Inflammation is one of the main determinants of post-stroke recovery. Emerging data suggest that inflammatory cells play complex and multiphasic roles after ischemic stroke, and most cell types display both beneficial and detrimental effects (1). There is growing evidence that inflammation in the early phase after ischemic stroke is deleterious, and the downregulation of inflammation in the acute phase of stroke is likely to be neuroprotective (1, 2). Attempts to translate anti-inflammatory agents in ischemic stroke have remained largely unsuccessful at reducing lesion progression (2).

Most of these interventions have been targeted at a specific cytokine, chemokine, or inflammatory process, even though the inflammatory cascade is multifaceted with a significant amount of inherent redundancy (3). Fasting has been shown to be a successful intervention to lower inflammation in different species through various mechanisms, including the downregulation of the P13K/Akt/mTOR signaling pathway (4). In rodent models, fasting has been shown to reduce inflammation in stroke (5) and cortical injury (6). However, most studies introduced the dietary intervention before the brain injury, which is - translationally speaking - irrelevant. This study introduces fasting after a focal model of cerebral ischemia in rats and investigates its acute effects on infarct volume, BBB breakdown, and inflammation at 24 hours.

## Methods

### Animals

All experimental procedures were approved by the UK Home Office (1986 Animal Act, Scientific Procedures), conducted in accordance with local ethical guidelines at the University of Oxford, and where possible, conformed to the ARRIVE and IMPROVE guidelines for animal and pre-clinical stroke work (7, 8). Further details can be found in the Supplemental Material.

### Intervention

Focal brain ischemia was induced by endothelin-1 (ET-1) injection into the area of the right striatum, as previously described (9).

### Treatment

Animals in the fasting group had no access to food for 24 hours from the point of suture closure, with ad libitum access to water (n=11). The control group had ad libitum access to food and water (n=10).

### Infarct volume

Cresyl violet was used to stain five brain sections per animal which were +1.28 mm, +0.6 mm, +0.12 mm, -0.12, and -0.48 mm away from the bregma, respectively. To obtain the lesion volume, infarct areas were multiplied by the distance between the brain. Lesion volume was then presented as a percentage of the contralateral hemisphere.

### Blood analysis

For full blood count analysis, 5 ml of whole blood was drawn directly out of the heart in EDTA tubes and analyzed on the same day (Laboratory Haematology, John Radcliffe Hospital, Oxford, OX3 9DU, UK).

## Results

### Fasting reduces infarct volume

There was a statistically significant difference in infarct volume between the groups (t-test; p=0.0071, fasting: 2.685 ± 0.7762; control: 7.114 ± 1.307) (**Figures 1 A, B**).

**Figure 1.**
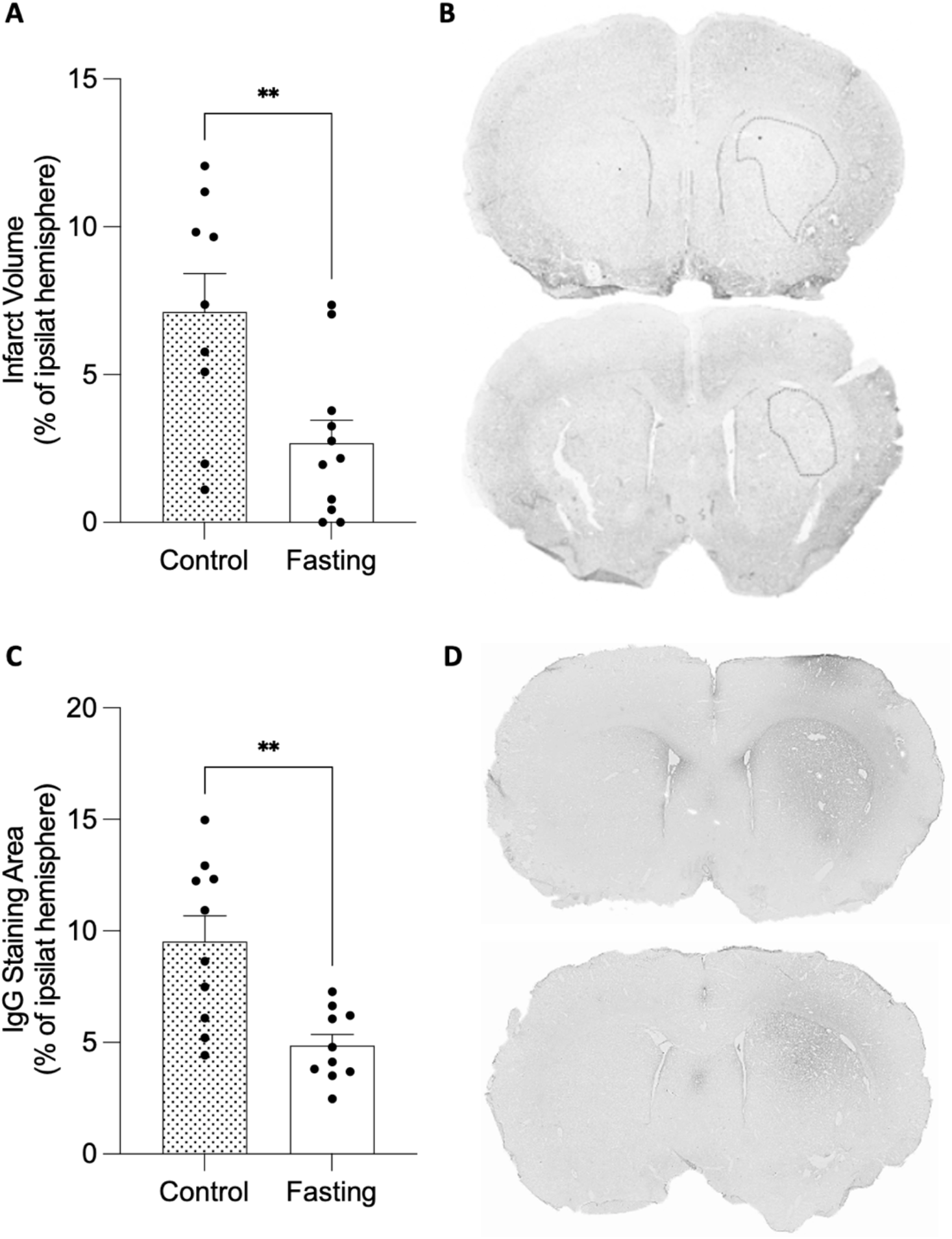
Fasting reduces infarct volume and BBB breakdown. **(A)** Infarct volume. **(C)** BBB breakdown visualized by IgG-positive staining. **(B, D)** Representative images of control and fasted animals. Data are mean ± SEM. **p<0.001. For infarct volume, n=9 for control and n=11 for fasted rats; for BBB breakdown, n=10 for both groups. Scale bar represents 1 mm.

### Fasting reduces BBB breakdown

BBB compromise was studied using IgG staining to visualize serum proteins in the CNS. IgG-positive staining was identified, and the IgG-positive area was then calculated as a percentage of the contralateral hemisphere. The mean value of BBB breakdown was significantly lower in fasted animals versus the control group (t-test, p=0.0016; fasting: 4.859 ± 0.5027; control: 9.523 ± 1.154) (**Figures 1 B, C**).

### Fasting does not affect CNS resident immune cells

Microglia and astrocytes (identified as Iba-1 and GFAP-positive cells, respectively) were quantified at 24 hours in an area of 1 mm^2^ within the cortex and striatum. As for microglial numbers in the striatum, there was no main effect of stroke (2-way ANOVA; p=0.0525), no main effect of treatment (p=0.6224), and no interaction between the effects (F(1,37)=1.976, p=0.1618). Šidák’s multiple comparisons tests found a significant increase in microglial numbers in control animals of the stroke compared to the sham group (**Figure 2 B**). In the cortex, no effect of stroke or treatment on microglial numbers was found (**Figure 2 A**). Similarly, no effect of stroke or treatment on astrocyte numbers was found in either the cortex or striatum (**Figure 2 C, D**).

**Figure 2.**
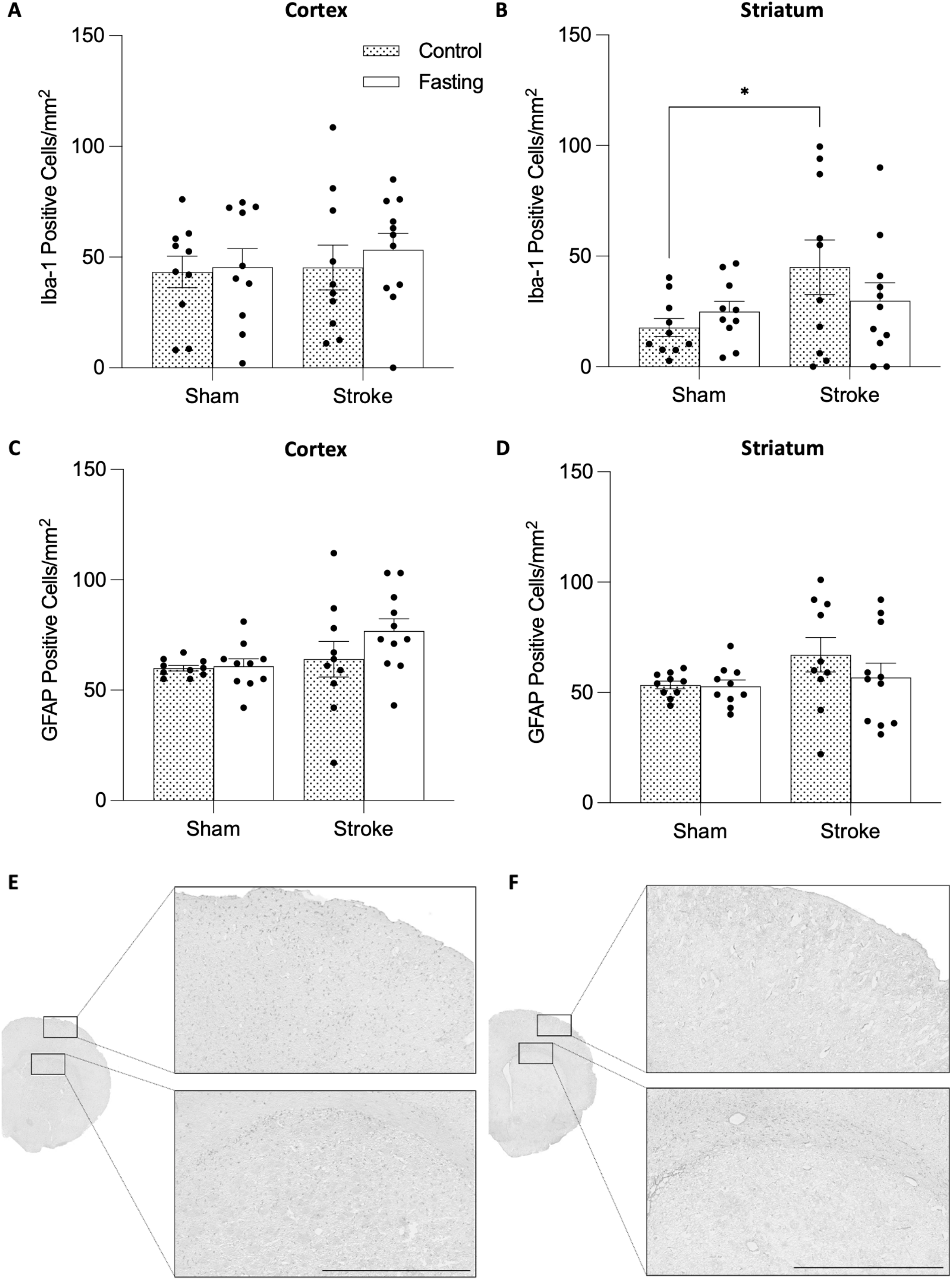
Fasting does not affect microglia or astrocyte numbers. Microglial numbers in the **(A)** cortex and the **(B)** striatum. Astrocyte numbers in the **(C)** cortex and the **(D)** striatum. **(E, F)** Representative images of microglia or astrocyte-staining, respectively, of the cortex and corpus callosum of a fasted rat. Data are mean ± SEM. *p<0.05. n=10/group for all groups. Scale bar represents 1 mm.

### Fasting reduces neutrophil infiltration into the striatum

The mean number of neutrophils in the CNS was calculated. There was a statistically significant difference between the two treatment groups (t-test; p=0.0249; fasting: 50.91 ± 16.01; control: 145.7 ± 36.92) (**Figure 3 A**).

**Figure 3.**
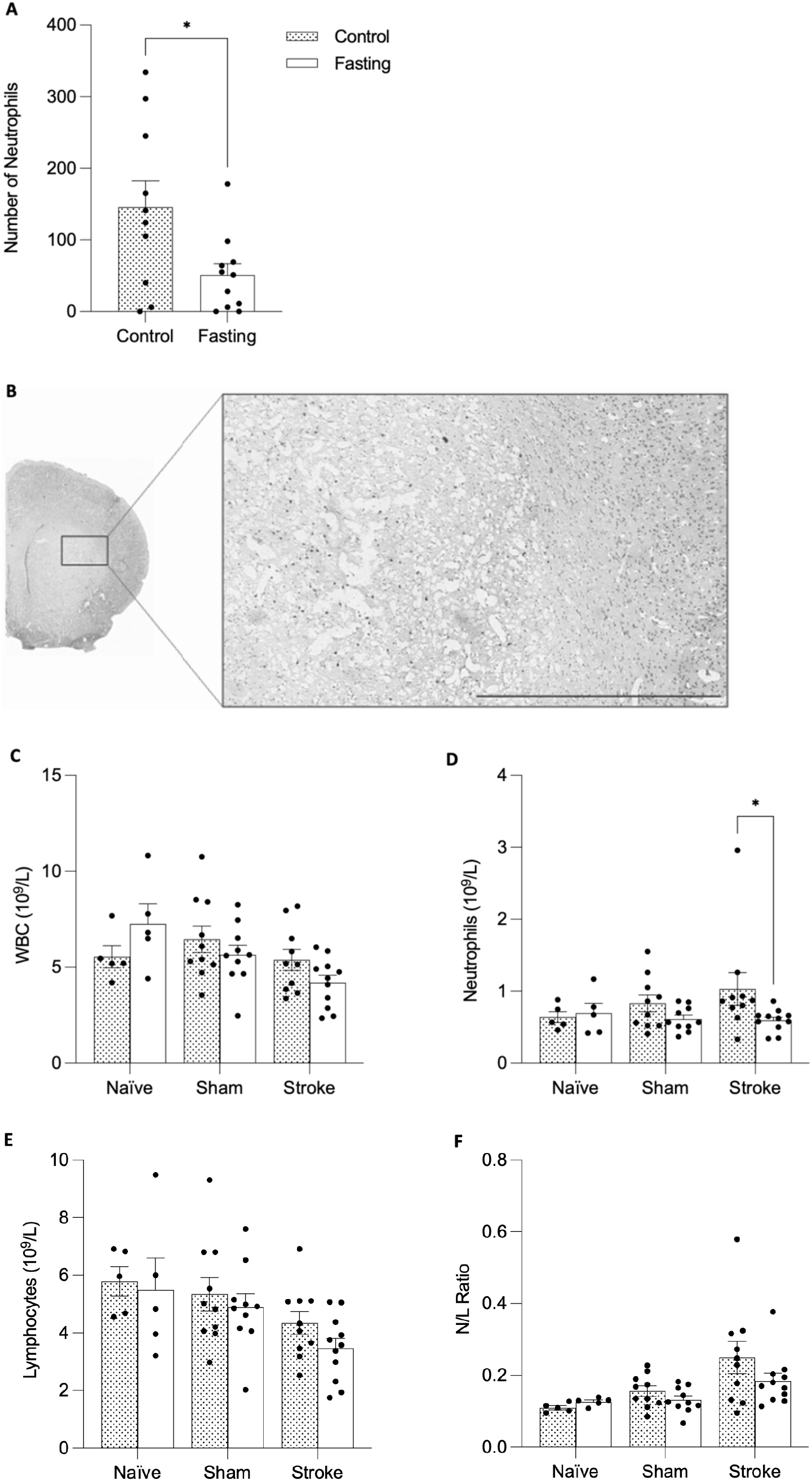
Fasting reduces central neutrophil infiltration and reduces blood neutrophils. **(A)** Infiltrating neutrophils in the striatum of post-ischemic rats. **(B)** Representative image of infiltrating neutrophils. **(C)** WBC, **(D)** neutrophils, **(E)** leukocytes, and **(F)** N/L ratio. Data are mean ± SEM. For histological analysis, n=10/group for all groups. For blood analysis, n=5 in naïve groups, n=10 in sham and stroke groups. *p<0.05. Scale bar represents 1 mm.

### Fasting reduces circulating neutrophils

At 24 hours post-stroke, a full blood count was performed, and neutrophil numbers, white blood cells (WBC), lymphocyte numbers, and the neutrophil/lymphocyte (N/L) ratio were analyzed. Considering neutrophils, there was no main effect of the surgery (2-way ANOVA; p=0.5823) or treatment (p=0.0889), and no interaction between the two (p=0.2611). Šidák’s multiple comparisons tests found that, in the stroke animals, fasting significantly reduced the circulating neutrophils (p=0.0400) (**Figure 3 D**). As for WBC, lymphocyte numbers, and N/L ratio, there was a main effect of the surgical intervention but no effect of treatment and no interaction between the two effects (**Figure 3 C, E, F**).

## Discussion

Unlike pharmacological treatments, fasting may be beneficial as it does not suffer from a significant side-effect profile (4, 10). However, the inherent biochemical and metabolic differences between rodents and humans must be considered when expanding and translating these studies. The dietary pattern of stroke patients is influenced by the clinical outcome, and dysphagia, for example, is a common problem in the clinic, but no controlled studies of its effects on stroke outcomes have been performed. In contrast, in rodents, there is pressure to make sure the animals are adequately eating, and often those that do not because of severe injury are excluded from the studies. This study sought to investigate the impact of a 24-hour water-only fast in rats after ET-1-induced cerebral ischemia. Fasting significantly reduces infarct volume and BBB breakdown and offsets the increased number of circulating neutrophils and infiltration of neutrophils into the striatum caused by ischemia. These results demonstrate that fasting is a promising way of mitigating stroke injury by potentially reducing inflammation and ameliorating the breakdown of the BBB.

Inflammation is an essential aspect of the post-stroke landscape, and fasting is known to be anti-inflammatory. Weston and colleagues investigated the temporal relationship between ischemic damage and neutrophil numbers in the ET-1 stroke model (11). They found that infarct volume and neutrophil infiltration into the brain peaked at three days and that neutrophil numbers positively correlated with the volume of infarcted tissue (11). In our study, we find that at 24 hours, alongside increased neutrophils in the blood, neutrophils have infiltrated the brain and that this infiltration is reduced by the post-ictal fasting period. By extending the endpoint to 72 hours, future studies may be able to establish whether this effect persists beyond the acute phase.

Data from the cardiac field suggests fasting would be beneficial (12), but would also benefit from comparison to more sustainable longer-term options such as caloric restriction or timed feeding (intermittent fasting). Given the fluid nature of the inflammatory response post-stroke, it’s even possible that altering the feeding strategy over the immediate post-stroke period would be beneficial to maintain the reduced inflammatory response - starting with fasting, followed by timed feeding, followed by caloric restriction. However, these investigations were outside the scope of this preliminary study.

The current study didn’t find an effect of fasting on the numbers of microglia and astrocytes at 24 hours. While microglia are known to react to hypoxic stimuli with a brief and local increase in number and change in morphology, astrocytes express more long-term and global astrogliosis (13, 14). The current study’s approach to quantifying, but not morphologically defining microglia is likely limiting our capacity to capture the subtle subacute changes in CNS inflammation.

In conclusion, this work sought to examine the effects of fasting on infarct volume, BBB integrity, and post-ischemic inflammation at 24 hours. The results presented are promising and suggest fasting as a potential adjunct treatment option in ischemic stroke. Whilst more data is needed to explore the effects of fasting in the chronic phases after injury, this initial dataset has yielded promising results for fasting to be used as an additional treatment option that does not bear the risk of pharmacological side effects, is low cost, and is widely accessible.

## Acknowledgments

The authors would like to acknowledge the assistance of Liam Silvera at the University of Caltech.

## Supplemental Material

Detailed Methods, Detailed Results, Supplementary Figure 1, Supplementary Table 1.

