## Supplemental Material for "The effects of fasting on ischemic infarcts in the rat"

**Detailed Methods**

*Animals:* Male Wistar Han rats (250-320 g, Envigo Research Model Services in Blackthorn, England) were housed in individually ventilated cages under a 12-hour light/12-hour dark cycle with ad libitum access to water. Before the intervention, access to food was unrestricted and strictly controlled or restricted in the first 24 hours after the stroke. The principal investigator began daily animal handling and weighing the animals three days before the surgery.

*Sufficient statistical power:* Preliminary data on the effects of fasting after stroke are limited. Power calculations were based on previous work from our group. A sample size of ten rats per group was used, which was thought to be sufficient to detect biologically significant effects of infarct volume as the primary outcome measure while minimizing the number of animals required to achieve the research objectives of this study.

*Controls:* Appropriate control groups were included in all experiments. Control animals had ad libitum access to food and water and did not receive any pharmacological treatment. To control for the stroke itself, sham animals with the respective treatment (n=10 for fasting, n=10 for control) were included. Sham surgery followed the identical protocol to the endothelin-1 (ET-1) model but with sterile saline injection rather than ET-1. Further, to control for the surgical intervention itself, naïve animals that did not undergo surgery were included (n=5 for fasting, n=5 for control). Throughout the chapter, the different study groups are referred to as “stroke”, “sham” and “naïve”, respectively. Pre-defined inclusion criteria were the presence of data on the infarct volume and BBB breakdown as the primary outcome measures, while pre-defined exclusion criteria were a post-procedural welfare score of at or above 7 (**Supplementary Table 1**) or death. No animal had to be excluded from this study. The primary outcome measures were infarct volume and BBB breakdown, while the secondary outcome measures were inflammatory markers in the brain and blood.

*Randomization and blinding:* To prevent accidental bias and confounding in the *in vivo* experiments, treatment allocation of fasting versus feeding was alternated between animals. Further, tissue was randomized after collection (YC) through changing animal identification numbers, and blinding was continued throughout the experiment wherever possible until data acquisition was complete.

*Welfare assessment:* Post-procedural monitoring included a welfare assessment performed every 30 minutes for the first 3 hours after the surgery and then every 12 hours, using the scoring system described in **Supplementary Table 1** and the recommendations of the IMPROVE guidelines (7).

*Endothelin surgery:* In this experiment, focal cerebral ischemia was induced using the ET-1, which was based on ethical considerations to using a stroke model that involves a less severe surgery that allows for subsequent fasting. During the surgery, the core body temperature of all animals was maintained at 37.0 ± 0.5°C using a rectal thermometer connected to a feedback-controlled heating pad (Harvard Apparatus, Cambourne, UK). Physical parameters, including body temperature and respiratory rate, were checked and recorded every 5 minutes throughout the surgical intervention. Respiration was kept between 50 and 60 breaths per minute by adjusting isoflurane concentration. Focal brain ischemia was induced by ET-1 injection into the area of the right striatum, similar to a previously described method (9). Briefly, the rat was deeply anesthetized with 5% isoflurane in 70% N_2_ and 30% O_2_ and maintained at 1-2% isoflurane throughout the intervention. After weighing, blood glucose and ketone bodies were measured before the head was shaved and disinfected using a 70% ethanol and 30% chlorhexidine solution. The animal was secured in a stereotaxic frame before a midline incision at the top of the head was made, and the needle navigated to the coordinates of the MCA territory in the striatum of the right hemisphere (AP +1.0, ML -3.0, SI -4.0 mm). A small hole was drilled in the skull, through which a needle was inserted, and 1 μl of ET-1 (25 pmol) or saline (for the control group) was slowly injected over 2 minutes. The head wounds were cleaned and closed (4-0 Vicryl Rapide Undyed 1x18” P-3, Somerville, US). Animals received 0.05 mg/kg marcaine into the wound to alleviate pain and 2 ml of saline solution subcutaneously. The rectal thermometer was removed, isoflurane was turned off, and the gases were changed to 0% N_2_O and 100% O_2_. Upon awakening, the animal was put in a pre-warmed cage, using a heating mat underneath. The animal was closely monitored for an additional 3 hours after awakening, with post-surgical welfare check-ups every 30 minutes.

*Tissue processing:* After 24 hours from the start of treatment, rats were deeply anesthetized with 5% isoflurane in 70% N_2_ and 30% O_2_. Rats were weighed, blood glucose and blood ketone bodies were measured (On Call GK Dual Blood Glucose & Ketone Monitoring System, Acon Laboratories, San Diego, CA, USA), and then killed by intraperitoneal pentobarbital injection (800 mg/kg). 5 ml of blood was drawn from the heart into EDTA tubes, and full blood count was analyzed on the same day (Laboratory Haematology, John Radcliffe Hospital, Oxford OX3 9DU, UK). Animals were transcardially perfused using heparinized saline and 4% PFA in PBS. Brains were collected, post-fixed in 4% PFA in PBS overnight, and changed into a 30% sucrose solution in PBS. For the naïve group, fresh non-perfused brains were collected and sliced into 2 mm thick coronal sections with an ice-cold stainless-steel matrix (Kent Scientific). 1 mm^2^ tissue samples of the striatum and cortex of both hemispheres were snap-frozen on dry ice.

*Immunohistochemistry:* Perfused brains were dehydrated through graded ethanol solutions and embedded in OCT mounting medium before snap-frozen on dry ice. Using a cryostat, 10 μm-thick serial sections in the coronal plane were collected 0.5 mm anterior to posterior of the lesion on gelatinized slides and stored at -80°C until further use. Brain sections were rehydrated through graded ethanol solutions. Nonspecific binding was blocked using 10% of serum of the species the secondary antibody was raised in (diluted in PBS) for 1 hour at RT and incubated in primary antibody in PBS at 4°C overnight. After that, sections were rinsed in PBS, and a secondary antibody was added at 1:500 in PBS for 45 minutes at RT. Antibody binding was visualized with 3,3’-diaminobenzidine. Sections were dehydrated through graded ethanol solutions and cleared with Histo-Clear II (National Diagnostics, HS2021GLL). The slides were mounted with glass coverslips using an anti-fade fluorescence mounting medium (Dako, USA) and imaged with a microscope scanner (Manual Whole Slide Imager 2017b-31, Olympus Life Science) with 10 x magnification.

*Statistical analysis:* Statistical analysis was carried out in Prism 6 (Graphpad, USA). For analysis of infarct volume, BBB breakdown, and neutrophil infiltration into the brain, an unpaired t-test was used. For weight, blood glucose and ketone bodies, microglia and astrocyte count, and full blood cell count, 2-way ANOVA was used with Dunnett’s multiple comparisons test. The D’Agostino and Pearson normality tests were performed on all data, and the appropriate statistical tests were chosen based on the normality of the data. α of 0.05 was considered statistically significant, and the results are presented as mean ± standard error (SEM). * p<0.05, ** p<0.01, *** p<0.001, **** p<0.0001.

**Detailed Results**

*Fasting decreases body weight:* Bodyweight measurements were recorded immediately before the surgical intervention and at 24 hours, marking the experiment's end. The aim was to determine whether fasting affects body weight in animals that have experienced a stroke. There was no main effect of the surgery (2-way ANOVA; p=0.3582), but a main effect of treatment (p<0.0001) and an interaction between the two effects (F(2, 45)=4.130, p=0.0225). Šidák’s multiple comparisons tests showed that fasting significantly reduced body weight in naïve, sham, and stroke animals (p<0.0001 for all) (**Supplementary Figure 1 A**).

*The effects of fasting on blood glucose levels and ketone bodies:* The next aim was to investigate whether fasting affects blood glucose and ketone levels in animals that have experienced a stroke. At 24 hours post-stroke, blood glucose and ketone body levels were measured. There was a main effect of the surgery (2-way ANOVA; p=0.0412) and treatment (p<0.0001), but no interaction between the two (p=0.0661). Šidák’s multiple comparisons tests showed significant decreases in glucose levels in fasted animals in all intervention groups (p=0.0115 for naïve, and p<0.0001 for sham and stroke animals) (**Supplementary Figure 1 B**). For ketone body levels, there was a main effect of the surgery (p=0.0631), a main effect of treatment (p<0.0001), and no interaction between the two (F(2,45)=2.262, p=0.1158). Šidák’s multiple comparisons tests showed that fasting significantly increased ketone bodies in all treatment groups (p=0.0105 for naïve, and p<0.0001 for sham and stroke animals) (**Supplementary Figure 1 C**).

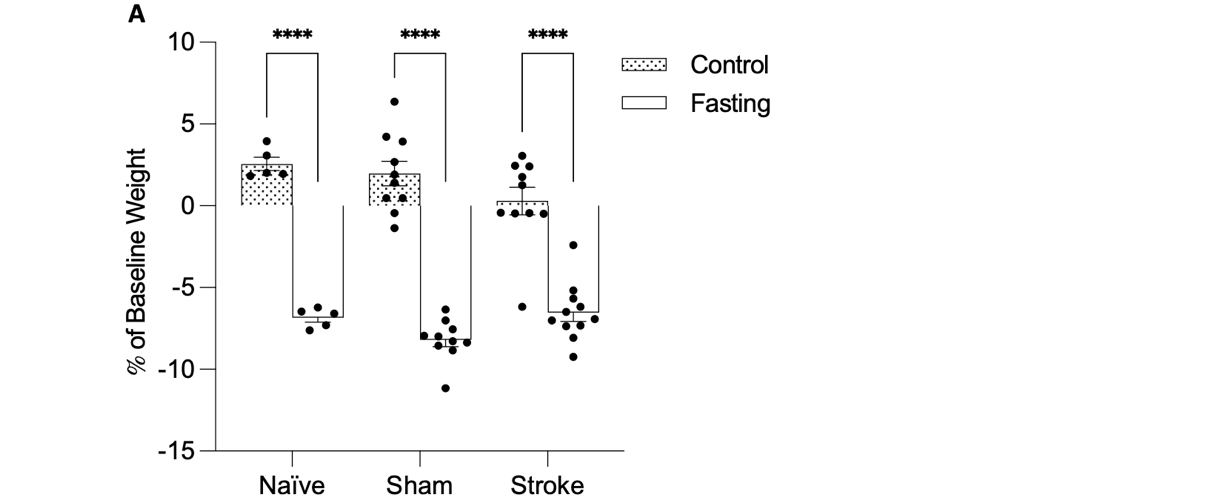

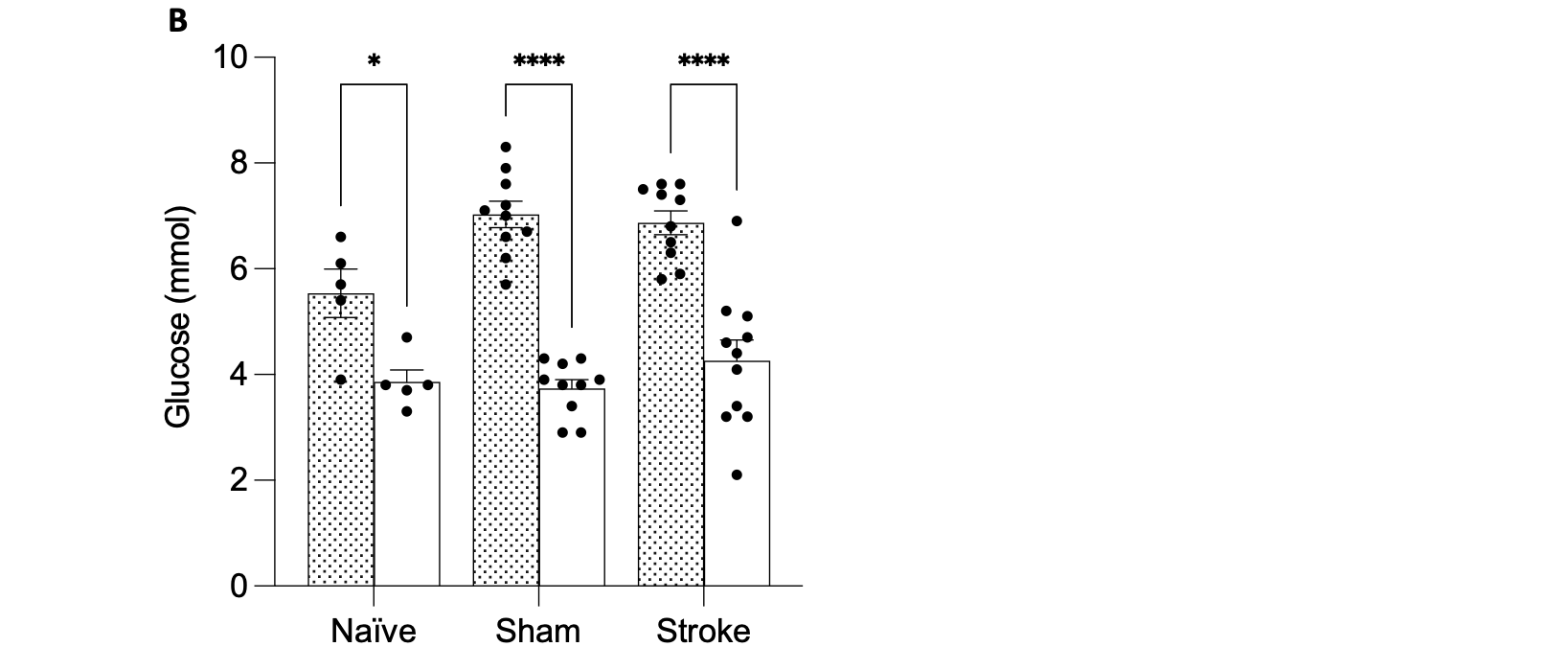

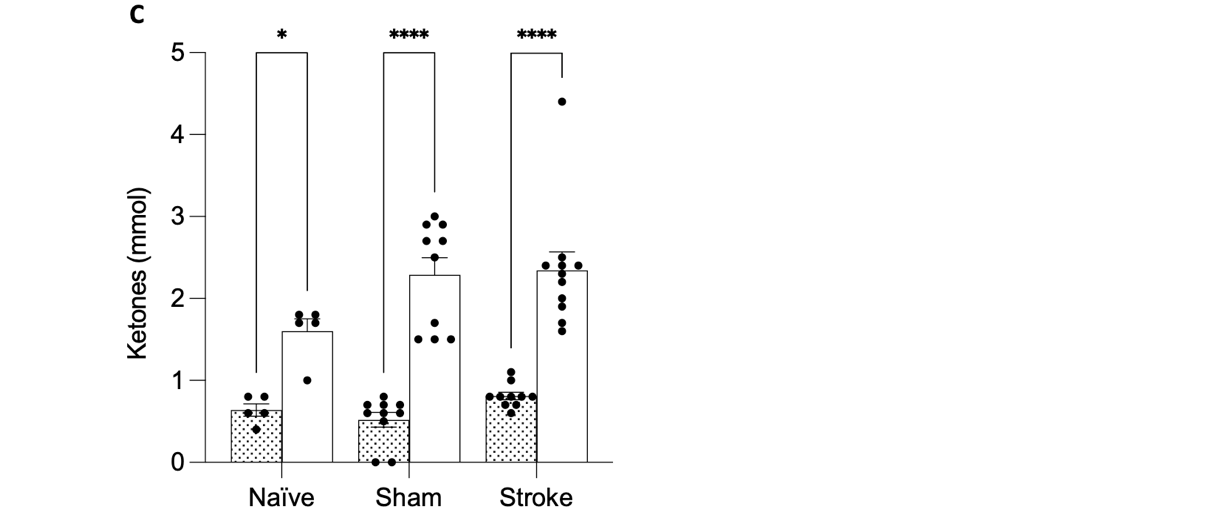

**Supplementary Figure 1.** Fasting reduces body weight and blood glucose levels and increases ketone bodies at 24 hours. **(A)** Body weight, **(B)** blood glucose, and **(C)** ketone bodies. Results are presented mean ± SEM. *p<0.05, ****p<0.0001. n=5 in naïve groups, n=10 in sham and stroke groups.

|  | Scoring |
| --- | --- |
| **Appearance** |  |
| Normal | 0 |
| Lack of grooming | 1 |
| Coat staring/ocular discharge/nasal discharge | 2 |
| Piloerection/hunched posture | 4 |
| **Food and water intake** |  |
| Maintaining body weight within 5% of baseline weight | 0 |
| Weight loss 5-10% | 1 |
| Weight loss 10-15% | 2 |
| Weight loss > 10-15% | 4 |
| Weight loss > 10-15%, sustained for ≥ 48 hours | 12 |
| **Natural behaviour** |  |
| Normal | 0 |
| Minor change in spontaneous activity | 1 |
| Substantial change in spontaneous activity | 2 |
| Restless or very still | 4 |
| Chewing limb, lameness, loss of body supply to leg | 12 |
| **Cerebral function** |  |
| Normal | 0 |
| Minor/moderate circling and/or hemiparesis | 2 |
| Severe circling and/or hemiparesis/hemiplegia | 6 |
| Fitting or ataxia | 12 |
| **Postoperative complications** |  |
| Difficulty in breathing | 4 |
| Excessive bleeding | 4 |
| Infection | 4 |
| Wound breakdown | 4 |

**Supplementary Table 1.** Welfare assessment ranging from 0 to 56. Scores of less than 4 were considered mild, whereas 7 and above were considered a severe impairment, respectively. A score of 7 and above were pre-defined humane endpoints.
